# VCP protects neurons from proteopathic seeding

**DOI:** 10.1101/2021.07.12.452081

**Authors:** Jiang Zhu, Sara Pittman, Dhruva Dhavale, Rachel French, Jessica N. Patterson, Salman Kaleelurrrahuman, Yuanzi Sun, William J. Buscher, Jan Bieschke, Paul Kotzbauer, Yuna Ayala, Albert A. Davis, Conrad Weihl

## Abstract

Uptake and spread of proteopathic seeds, such as αS, Tau, and TDP-43, contribute to neurodegeneration. The cellular machinery necessary for this process is poorly understood. Using a genome-wide CRISPR-Cas9 screen, we identified Valosin Containing Protein (VCP) as a suppressor of αS seeding. Dominant mutations in VCP cause multisystem proteinopathy (MSP) with muscle and neuronal degeneration. VCP inhibition or disease mutations increase αS seeding in cells and neurons. This is not associated with an increase in seed uptake and is similar to treatment with the lysosomal damaging agent, LLoME. Intrastriatal injection of αS seeds into VCP disease mice enhances seeding efficiency compared with controls. This is not specific to αS since VCP inhibition or disease mutations increased TDP-43 seeding in neurons. These data support that VCP protects against proteopathic spread of pathogenic aggregates. The spread of distinct aggregate species may dictate pleiotropic phenotypes and pathologies in VCP associated MSP.

## INTRODUCTION

α-synuclein (αS) is the principal component of protein inclusions that are found in a family of neurodegenerative disorders known as synucleinopathies ^1^. Synucleinopathies include Parkinson’s Disease, Diffuse Lewy body disease (DLB), multiple system atrophy and REM sleep behavior disorder (RBD) ^1^. αS contains an amyloid-like region making it prone to aggregate. Several lines of evidence suggest that aggregated αS can seed the fibrillization of soluble, monomeric αS ^2^. This process may relate to disease pathogenesis and progression ^3^. αS fibrils enter neurons via endocytosis and template new αS aggregates within the cytoplasm ^4^. This leads to synapse loss, neurodegeneration, and ultimately the release of αS fibrils to adjacent cells resulting in aggregate propagation along interconnected neurons ^3^. The cellular process of proteopathic seeding consists of several regulated steps. These include seed uptake, vesicular trafficking, endolysosomal escape and templated conversion of cytosolic αS ^5^.

One route of seed uptake is endocytosis. αS seeds can enter cells and neurons via the endocytic system. These seeds then penetrate the endolysosomal membrane facilitating their escape into the cytoplasm ^5^. Pharmacologically blocking cellular uptake or increasing endolysosomal membrane damage can decrease and increase seeding efficiency respectively ^5^. Previous studies have identified genetic modifiers of αS toxicity associated with its intracellular expression in yeast or *C. elegans* ^6^. However, genetic modifiers of proteopathic seeding have not been explored.

VCP (also called p97, or cdc48 in yeast) is a ubiquitin-directed AAA-ATPase implicated in multiple forms of neurodegeneration ^7^. Dominantly inherited mutations in VCP cause multi-system proteinopathy (MSP) which is associated with multiple variably penetrant phenotypes that include inclusion body myopathy, frontotemporal dementia, ALS, and Parkinsonism ^7^. Just as the phenotypes are variable, VCP patients develop varied aggregate pathologies that include αS, TDP-43, Tau, SQSTM1 and ubiquitin inclusions ^8–11^. How VCP disease mutations lead to cellular degeneration and protein inclusions is unclear. VCP affects the trafficking and clearance of polyQ aggregates in vitro ^12^. VCP is also necessary for both autophagic and proteasomal degradation of ubiqutinated proteins including TDP-43 and ER associated proteins via ERAD ^7^. VCP has more recently been proposed to behave as a protein disaggregase specifically acting upon pathologic tau aggregates ^11^. By binding to distinct adaptors, VCP alters it functionality allowing it to participant in its many other functions such as ERAD, vesicular trafficking, DNA repair, and cell cycle regulation ^7^.

VCP disease mutations alter its association with distinct adaptors ^13, 14^. Specifically, VCP disease mutations have reduced binding to UBXD1 and increased interactions with Ufd1/Npl4 creating both a loss and gain of function with regard to UBXD1 and Ufd1 dependent processes ^13–16^. Notably, a VCP-UBXD1 dependent complex is recruited to damaged endolysosomes ^15^. This complex recruits the deubiquitinase YOD1 which cleaves K48 linked ubiquitin chains from the lysosomal membrane facilitating lysophagic degradation ^15^. VCP inhibition, loss of UBXD1, or VCP disease mutant expression lead to a delay in the clearance of damaged late endosomes resulting in the accumulation of galectin-3 positive puncta in both VCP mouse models and patient tissue ^15, 17^.

MSP patients are pathological characterized as a TDP-43 proteinopathy ^18^. MSP patient tissue accumulates aggregated and insoluble TDP-43 in affected muscle and CNS tissue ^18^. 90% of MSP patients have myopathy that precedes dementia by ~10 years ^19^. Whether TDP-43 aggregate pathology spreads from muscle to motor neuron and ultimately the cortex is not known. TDP-43 contains an intrinsically disordered or prion-like domain that facilitates its templated aggregate conversion ^20^. Like αS, TDP-43 aggregates can serve as proteopathic seeds that propagate in cell and mouse models ^21, 22^.

Functional genomic screens are a powerful tool to identify proteins participating in distinct cellular pathways. In this study, we utilized an αS seeding FRET biosensor to screen a CRISPR knockout library. This approach identified multiple suppressors of αS seeding of which the AAA ATPase, VCP, was further explored both *in vitro* and *in vivo*.

## RESULTS

### Genome-wide CRISPR knockout screen identifies genes protective against αS seeding

To identify genes that regulate αS seeding, we performed a genome-wide screen in a previously described HEK293 αS CFP/YFP biosensor cell line (αS biosensor) (Figure 1A) ^4^. Exogenously applied pre-formed fibrillar αS (αS PFF), but not monomeric αS, seeds the aggregation of soluble intracellular αS resulting in a quantifiable change in FRET efficiency. We clonally expressed spCAS9 in αS biosensors and then infected with a pooled Brunello gRNA library covering 19114 different genes/ 4 gRNA each and 1000 non-targeting controls at a low MOI (<0.3) for seven days with puromycin selection. The pooled knockdown αS biosensor maintained its normal αS seeding capacity (Figure S1). These biosensors were treated with αS PFF as previously described and flow-sorted into FRET positive and FRET negative groups. DNA was isolated from FRET positive, FRET negative, and the unseeded total cell population. Then next-generation sequencing was performed to identify the guide RNAs (gene knockdowns) represented in each group. This sequence data was analyzed via the MegaCK algorithm in both the FRET positive and negative populations compared with untreated control. 110 genes were enriched in the FRET positive population as compared to total population, and 43 genes were underrepresented in the FRET negative group versus total population (FDR<0.05 and fold change >2 or <0.5) (Figure 1B). These 153 genes were considered “protective” or suppressors of αS seeding in the biosensor line (Figure 1B-C; Table S1).

**Figure 1:**
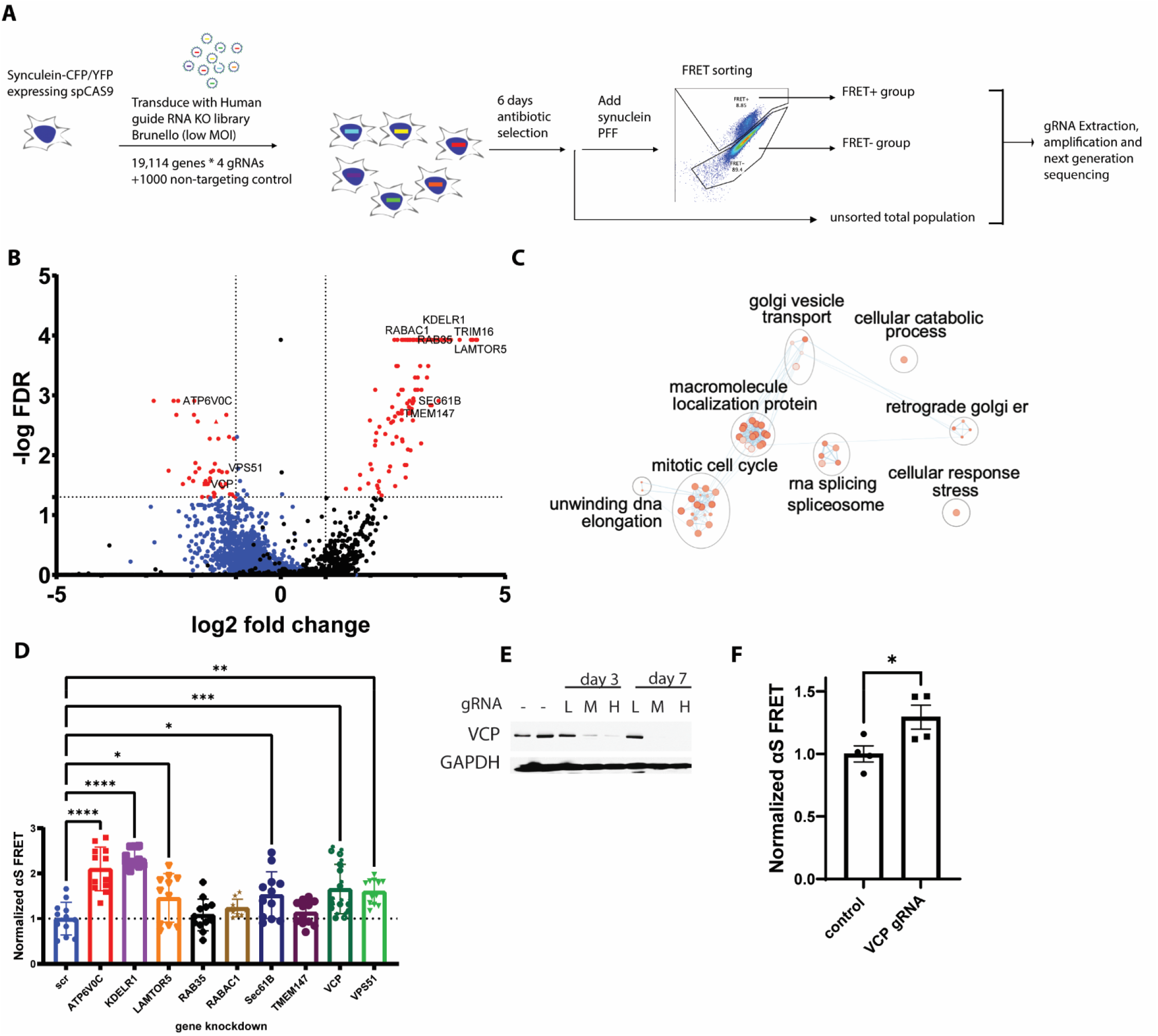
Genome-wide CRISPR/Cas9 screen identifies genes protective to αS seeding. (A) αS biosensor line stably expressing spCAS9 was transduced with sgRNA lentiviral library (Brunello). Following antibiotic selection, biosensors were seeded with αS PFF and flow sorted 24 hours later. Genomic DNA from positive and negative groups as well as unsorted total population were collected and decoded by NGS. (B) Volcano plot of genes identified in the screen. Colored in blue are all the genes plotted from FRET-group, while colored in black are those plotted from FRET+. Red dots are protective hits from both groups (5% FDR). Those genes with a fold change <0.5 were underrepresented in FRET-cells and genes >2 were overrepresented in the FRET+ cells. (C) Pathway analysis of 153 protective genes via g:profiler. The enriched pathway is visualized by cytoscape. (D) Normalized FRET of siRNA knockdown and αS PFF treated biosensors using 9 protective genes identified in the screen. (n ≥9 repeats; *** p < 0.001, ** p < 0.01 and * p < 0.05 by one-way ANOVA. error bars are ± S.E.M.) (E) Immunoblot with VCP and GAPDH of cell lysates from αS biosensors/spCas9 cells treated with a VCP gRNA (F) Normalized FRET of cells in (E) treated with 24-hour αS PFF. (P = 0.0439 by a two-tailed t-test). n = 4 biologically independent FRET assay. Data is mean ± S.E.M.

Our screen identified genes and pathways previously identified in other screens for αS toxicity. Notably, we identified 15 genes associated with ER-Golgi-endosome trafficking. These included VPS51 and VPS52 which are components of the Golgi-associated retrograde protein (GARP) complex. The GARP complex interacts with PD associated protein, LRRK2, and deletion of either VPS51 or VPS52 homologs in yeast, increases αS accumulation and toxicity ^23^. Other modifiers not previously identified in screens include ATP6V0B, ATP6V0C, and ATP6V1A that encode subunits of Vacuolar-type ATPase (V-ATPase). ATP6V0B KD inhibits autophagic degradation and increases αS aggregation ^24^ and is downregulated in patients with αS inclusions ^25^. Pathway analysis identified an enrichment in genes associated with the cellular stress response such as VCP, SEC61B and KDELR1. Notably the ER stress response is upregulated in PD brains and correlates with αS toxicity in multiple model systems ^26^.

We validated nine candidate suppressors (ATP6V0C, KDELR1, LAMTOR5, RAB35, RABAC1, SEC61B, TMEM147, VCP and VPS51) using siRNA knockdown in αS biosensors. Following 48 hours of siRNA treatment, αS PFF was added with Lipofectamine, and FRET efficiency was measured 24 hours later. Six candidate suppressors, when knocked down, increased FRET efficiency and included ATP6V0C, VPS51, KDELR1, SEC61B, LAMTOR5 and VCP (Figure 1D). To further confirm our findings with VCP, we generated a lentiviral vector expressing a VCP specific gRNA or control, infected spCAS9 αS biosensors for 7 days and treated with αS PFF. Similar to that seen with VCP siRNA, FRET efficiency was increased in VCP CRISPR KO αS biosensors (Figure 1E-F).

### VCP inhibition increases α-synuclein seeding efficiency

As previously reported, αS seeding as measured by FRET is not seen with the application of monomeric αS ^4^. Moreover, Lipofectamine is necessary for efficient FRET. The application of “naked” αS PFF to αS biosensors fails to increase FRET after 24 and 48 hours suggesting that Lipofectamine facilitates entry into the endolysosomal pathway ^4^. Consistent with this, the enhanced seeding efficiency and FRET signal following VCP siRNA treatment required the application of αS PFF with Lipofectamine and did not occur with monomeric αS or when αS PFF were added without Lipofectamine (Figure 2A).

**Figure 2:**
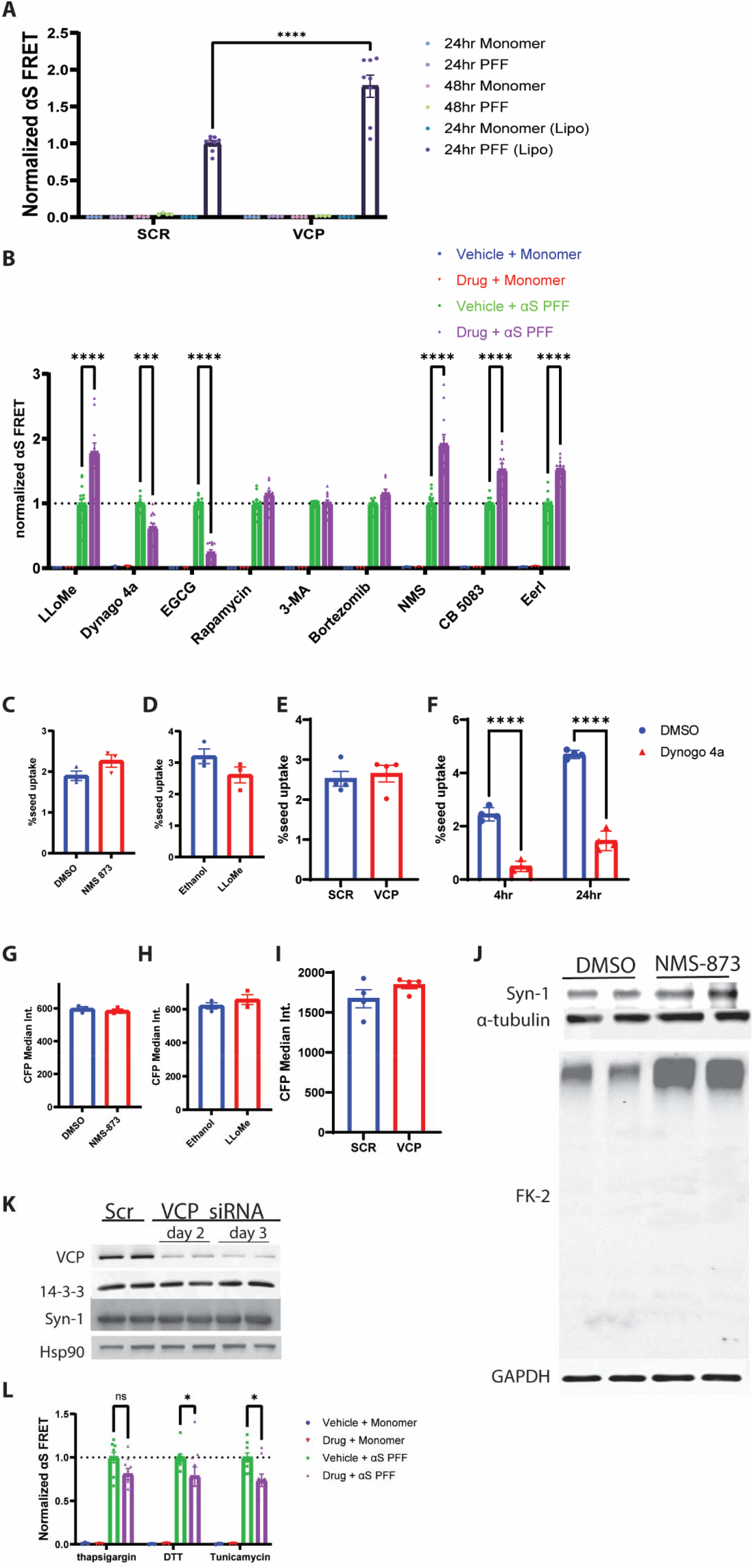
VCP inhibition enhances αS seeding independent of uptake or changes in αS protein levels. (A) αS biosensors were treated with scrambled control siRNA or VCP siRNA for 48 hours prior the application of αS monomer or PFF with or without Lipofectamine and measured for FRET efficiency at 24 or 48 hours. (B) Normalized FRET of αS biosensors co-treated with αS monomer or PFF with the indicated chemical compound or vehicle for four hours followed by washout. In the case of CB-5083 and Eer1, αS biosensors were treated for 24 hours. (n ≥9 repeats for each PFF treated conditions; n.s. for all monomer treated pairs; **** p < 0.0001 and *** p < 0.001 by two-way ANOVA in some PFF treated pairs as indicated. error bars are ± S.E.M.) (C) 647 fluorescent intensity (seed uptake) from αS biosensors co-treated with Alexa-647 tagged αS PFF and vehicle or LLoME for 4 hours. n=3 biological repeat. Data is mean ± s.e.m. (D) 647 fluorescent intensity (seed uptake) from αS biosensors co-treated with Alexa-647 tagged αS PFF and vehicle or NMS-873 for 4 hours. n=3 biological repeat. Data is mean ± s.e.m. (E) 647 fluorescent intensity (seed uptake) of αS biosensors treated with scrambled or VCP siRNA for 48 hours and then Alexa-647 tagged αS PFF for four hours. n=4 biological repeat. Data is mean ± s.e.m. (F) 647 fluorescent intensity (seed uptake) from αS biosensors co-treated with Alexa-647 tagged αS PFF and Dynogo-4a or DMSO for four hours and measured after 4 or 24 hours. n=4 biological repeat. Data is mean ± s.e.m. (G) CFP fluorescent intensity in αS biosensors following vehicle or LLoME for 24 hours. n=3 biological repeat. Data is mean ± s.e.m (H) CFP fluorescent intensity in αS biosensors following vehicle or NMS-873 for 24 hours. n=3 biological repeat. Data is mean ± s.e.m (I) CFP fluorescent intensity in αS biosensors following scrambled or VCP siRNA treatment for 48 hours. n=4 biological repeat. Data is mean ± s.e.m (J) Immunoblot of total α-synuclein (syn-1), FK-2, tubulin or GAPDH from αS biosensors treated with DMSO or VCP inhibitor NMS-873 for 4 hours. (K) Immunoblot of VCP, 14-3-3, total α-synuclein (syn-1) or HSP90 from αS biosensors treated with scrambled or VCP siRNA for 48 and 72 hours. (L) Normalized FRET of αS biosensors co-treated with αS monomer or PFF with the indicated chemical compound or vehicle for 24 hours. (n =9 repeats for each PFF treated conditions; n.s. for all monomer treated pairs; adjusted p = 0.0470 and p=0.0124 for DTT and Tunicamycin by two-way ANOVA. Error bars are ± S.E.M.)

In order to probe the role of VCP specifically at the time of endocytic entry into the cytosol, we modified our seeding protocol to a four-hour application with lipofectamine and subsequent washout of αS PFF from the media and determined this to be sufficient to seed αS aggregation as measured by FRET in a concentration-dependent manner at 24 hours (Figure S2A). Consistent with αS PFF entry through the endolysosomal pathway, treatment of αS biosensors with Dynogo-4a, a dynamin I/II inhibitor with αS PFF decreased FRET. Whereas a four-hour treatment at the time of αS PFF application with the lysosomal permeabilizing agent (LLoMe) significantly increased FRET (Figure 2B). Treatment of αS biosensors with the VCP inhibitor NMS-873 for four hours at the time of αS PFF application similarly increased seeding efficiency as measured by FRET (Figure 2B). The application of two additional VCP inhibitors, CB-5083, and Eeyarestatin I (Eer1) during αS PFF seeding also increased seeding efficiency measured by FRET (Figure 2B). This effect was αS PFF dependent since the application of VCP inhibitors or LLoMe in the presence of αS monomer did not increase FRET (Figure 2B). A similar experiment adding the polyphenol (−)-epigallocatechin gallate (EGCG), an anti-amyloid agent at the time of αS PFF application, resulted in a decrease in seeding efficiency (Figure 2B). Finally, the application of the proteasome inhibitor bortezomib or autophagy modulators (Rapamycin and 3-methyladenine (3-MA)) did not affect seeding efficiency (Figure 2B).

To see whether the increased seeding efficiency with VCP inhibition was due to an increase in αS PFF uptake, we employed a fluorescently conjugated αS PFF (αS-PFF 647). αS-PFF 647 retains seeding capacity in αS biosensors (Figure S2B). A four-hour αS biosensor treatment with αS-PFF 647 in the presence of LLoME or NMS-873 did not increase the amount of internalized αS PFF 647 (Figure 2C-D). Uptake was also unchanged when comparing scrambled and VCP siRNA KD (Figure 2E). In contrast, the application of Dynogo-4a significantly decreased αS PFF 647 uptake (Figure 2F). In addition, the increased seeding efficiency with VCP chemical inhibition or VCP siRNA knockdown was not due to an increase in the steady-state levels of αS as determined by immunoblot and αS fluorescence intensity via flow cytometry (Fig 2G-K). VCP inhibition and knockdown induce ER stress and activate the unfolded protein response (UPR) ^27, 28^. Treatment of αS biosensors with the ER stress inducing agents dithiothreitol (DTT), thapsigargin and tunicamycin for 24 hours following the application of αS PFF decreased FRET efficiency as compared to vehicle controls suggesting that the effect of VCP inhibition on seeding was ER stress independent (Figure 2L).

To explore the role of VCP in a more relevant system of αS seeding that does not require the use of the carrier Lipofectamine, we adapted a previously described assay which adds αS PFF to primary cultured hippocampal neurons (HNs) ^29^. The addition of αS PFF to the media of HNs for four hours, followed by media exchange, resulted in detergent-insoluble, high molecular weight αS, and phospho-αS as measured by fractionation immunoblot or via immunofluorescence using a phospho-αS antibody in HNs after five days (Figure 3A-B). The co-application of LLoMe and αS PFF for 4 hours followed by washout further increased the amount of phospho-αS as compared with the application of αS PFF and vehicle after five days (Figure 3B-C). These results were αS PFF dependent since monomeric αS failed to generate phospho-αS staining (Figure S3A). We performed a similar assay and treated HNs for four hours with the reversible VCP inhibitor ML240 and αS PFF. Notably, the application of ML240 for four hours at the time of αS PFF application increased the level of phospho-αS as compared with vehicle-treated control, while drug treated HNs with αS monomers show no phospho-αS (Figure 3D-E, S3A). Treatment of primary HNs with two different shRNAs against VCP for four days before seed application further demonstrated an increase in an αS PFF-dependent increase in phospho-αS staining compared with scrambled shRNA control (Figure 3F-G; S3B).

**Figure 3:**
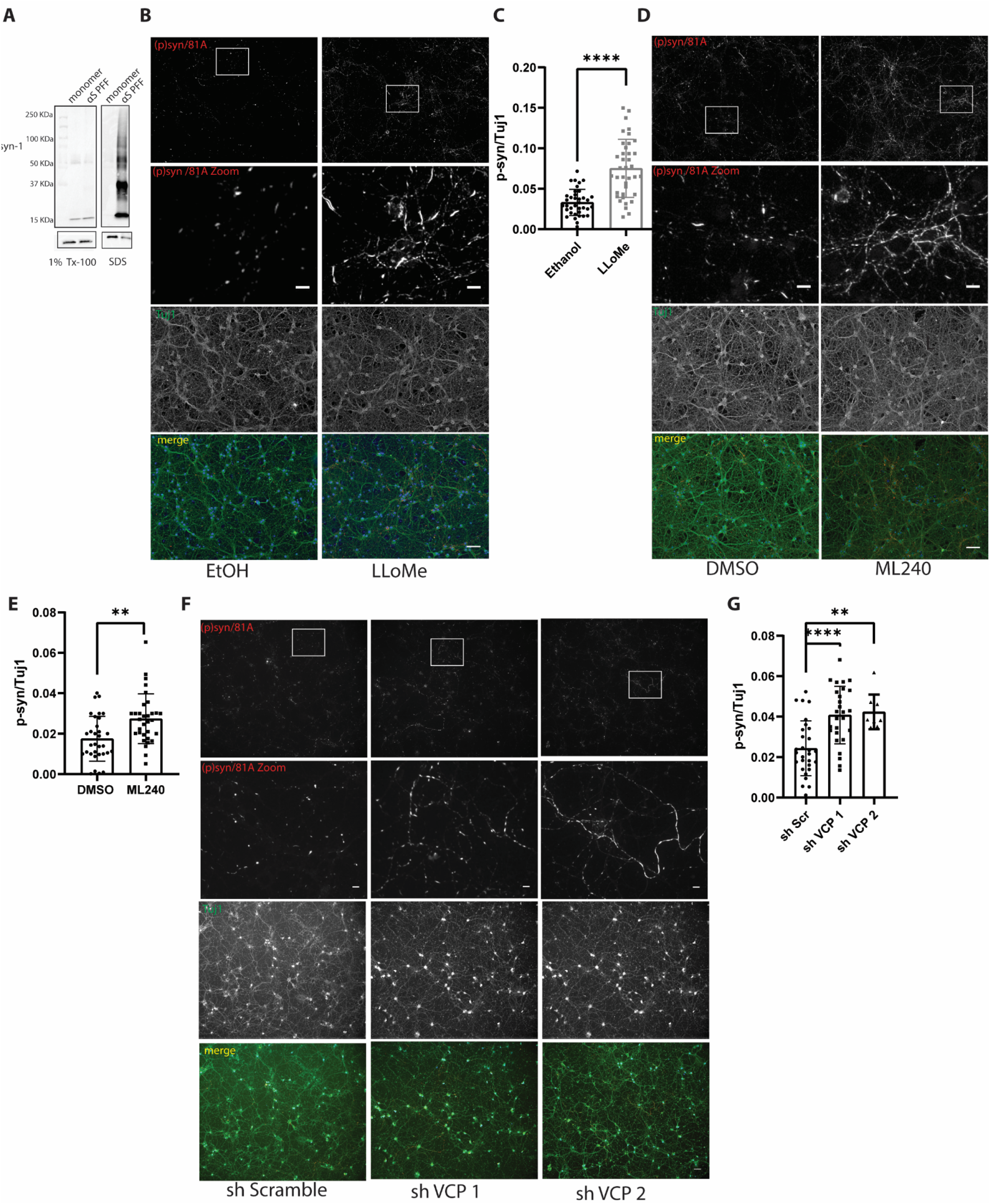
VCP inhibition enhances αS seeding in neurons. (A) Immunoblot for αS from detergent soluble and insoluble lysates of HNs treated for 4 hours with αS monomer or PFF and then harvested 5 days later. Note that the 2%SDS insoluble fraction has high molecular weight αS positive multimers. GAPDH is the loading control. (B) Immunofluorescence for phospho-αS and Tuj1 (neurite marker) in HNs co-treated with αS PFF and LLoMe (1uM) or vehicle for 4 hours. Immunostaining performed after 5 days. Scale bar =50 and 10μm for original and zoom-in figure respectively (C) Quantitation of phospho-αS/Tuj1 staining as in 3B (**** Multiple fields were captured in each conditions and quantified independently, n=37, 38 for ethanol and LLoMe groups respectively. Experiments were repeated from 3 different cultures. ****p < 0.0001 by student’s t-test. Error bars are ± S.E.M.). (D) Immunofluorescence for phospho-αS and Tuj1 in HNs co-treated with αS PFF and ML240 (100nM) or DMSO control for 4 hours. Scale bar =50 and 10μm for original and zoom-in figure respectively (E) Quantitation of phospho-αS/Tuj1 staining as in 3D (Multiple fields were captured in each conditions and quantified independently, n=32 for DMSO and ML240 group. Experiments were repeated from 3 different cultures. **p < 0.01 by student’s t-test. Error bars are ± S.E.M.). (F) Immunofluorescence for phospho-αS and Tuj1 in HNs infected with lentiviral vectors expressing scrambled or one of two different shRNAs targeting VCP for 2 days and then treated with αS PFF for 5 days. Scale bar =50 and 10μm for original and zoom-in figure respectively (G) Quantitation of phospho-αS/Tuj1 staining as in 3F (Multiple fields were captured in each conditions and quantified independently, n=29, 27 and 9 for scramble, VCP shRNA groups. Experiments were repeated from 3 different cultures. **p < 0.01, ***p < 0.001 by one-way ANOVA. Error bars are ± S.E.M.).

### VCP disease mutation expression increases αS seeding

VCP disease mutations affect a subset of VCP dependent cellular processes such as endocytic trafficking, nutrient sensing, autophagosome maturation, and, more recently, lysophagy ^7, 15^. This is due to an impairment in VCP mutant association with the adaptor UBXD1. We performed siRNA knockdown of VCP and the VCP adaptors UFD1, NPL4, UBXD1, and PLAA and the autophagy proteins ATG5 and SQSTM1 along with a scrambled control in αS biosensors. Following 48 hours of knockdown, αS biosensors were treated with αS PFF, and FRET was measured 24 hours later. Knockdown of VCP or its adaptor UBXD1 significantly increased FRET efficiency above controls (Figure 4A and S4A). shRNA knockdown of UBXD1 in primary HNs 4 days prior to αS PFF application also increased phospho-αS staining as compared with scrambled shRNA control (Figure 4B-C). To understand if VCP disease mutant expression increased αS seeding, similar to VCP and UBXD1 knockdown, we transfected αS biosensors with mCherry-tagged VCP-WT or one of three different VCP disease mutations (R95G, R155H, and A232E) for 24 hours. Treatment with αS PFF and quantitation of FRET efficiency 24 hours later in mCherry positive cells demonstrated an increase in FRET in VCP disease mutant expressing cells compared with VCP-WT control. Whereas cells not expressing mCherry did not show differences (Figure 4D and S4C).

**Figure 4:**
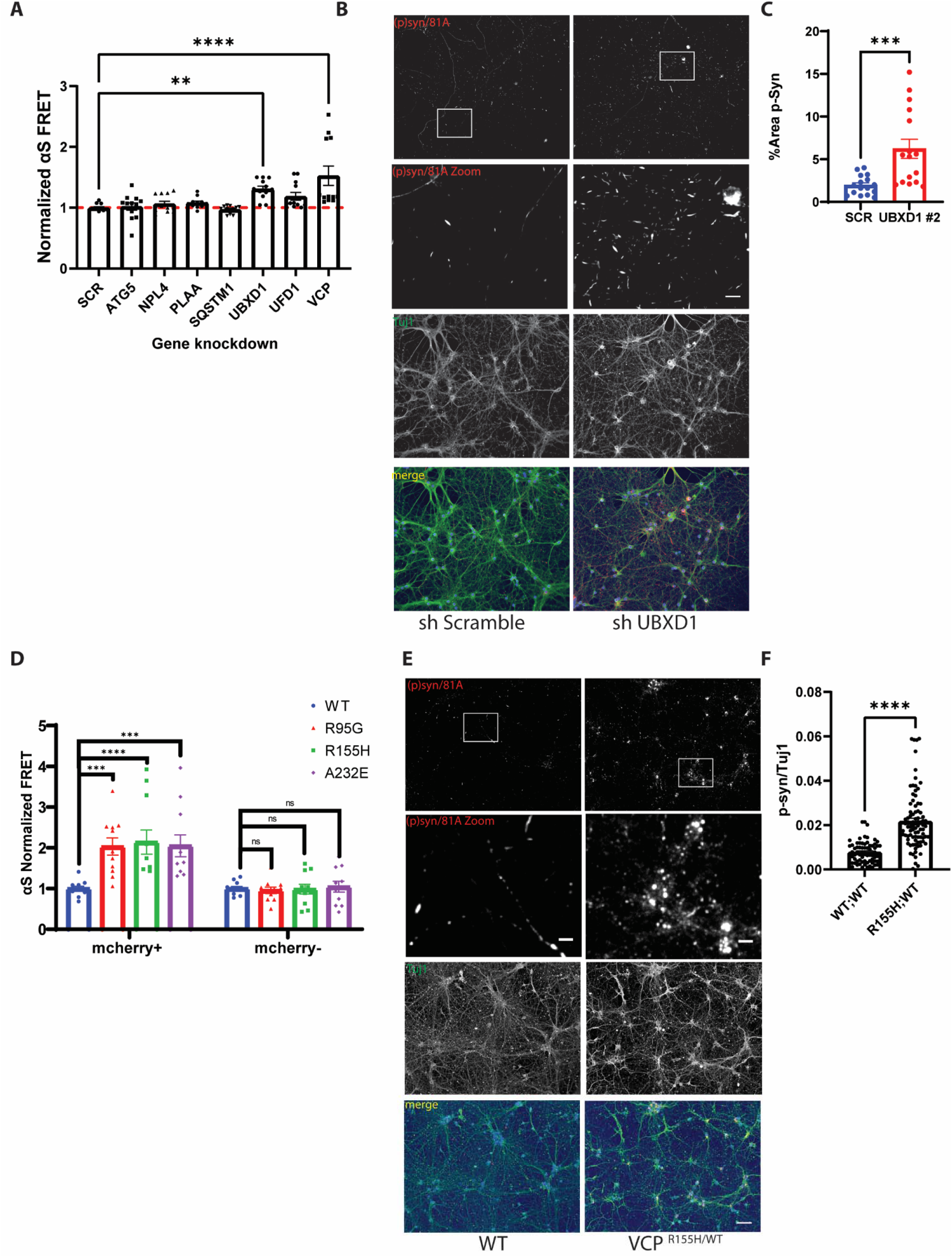
VCP disease mutations increase αS seeding. (A) αS biosensors were treated with scrambled control siRNA, siRNAs targeting the indicated genes for 48 hours prior the application of αS PFF and harvested for FRET efficiency at 24 hours. n ≥11 repeats for each KD. Adjusted p =0.0147 and p<0.0001 for UBXD1 and VCP KD respectively by one-way ANOVA. Error bars are ± S.E.M.) (B) Immunofluorescence for phospho-αS and Tuj1 in HNs infected with lentiviral vectors expressing scrambled or a shRNA targeting UBXD1 for 5 days and then treated with αS PFF for 5 days. (C) Quantitation of phospho-αS/Tuj1 staining as in 2B (*** P<0.0001). (D) αS biosensors were transfected with plasmids expressing VCP-WT, or one of three disease mutations (R95G, R155H and A232E) fused to an mcherry tag for 24 hours and then treated with αS PFF. FRET efficiency is quantified in mCherry+ and mCherry-cells separately and all normalized by VCP^WT^. (n=11 repeats for each group. ***p < 0.001; ****p < 0.0001; ns, no significance; two-way ANOVA with Dunnett’s correction) (E) Immunofluorescence for phospho-αS and Tuj1 in HNs form wild-type mice or mice carrying a VCP-R155H knockin allele (VCP^R155H/WT^) treated with αS PFF for 5 days. (F) Quantitation of phospho-αS/Tuj1 staining as in 4C (Multiple fields were captured in each conditions and quantified independently. Neurons coming from 7 and 12 independent cultures from WT and VCP^R155H/WT^ embryos. Outlier is removed by ROUT method, Q=1%, followed by Student’s t test. p<0.0001)

VCP-R155H mutation knockin mice have been previously generated and characterized ^30^. We cultured primary HNs from VCP^WT/WT^ and VCP^RH/WT^ embryos, treated them with αS PFF and then immunostained for phospho-αS five days later. VCP^RH/WT^ had a significant increase in phospho-αS staining compared with VCP^WT/WT^ controls (Figure 4E-F).

Using the same VCP-R155H mutation knockin mice, we examined the effect of pathogenic VCP mutations on αS seeding *in vivo*. VCP^RH/WT^ mice display no neuronal loss, TDP-43 inclusions, or pathologic features consistent with autophago-lysosomal dysfunction up to 13 months old (paper in print). To explore an additional VCP mouse model that only expresses a VCP disease mutant allele, we also used a mouse line that deletes the VCP-WT allele and only allows expression of a single VCP-R155C mutant allele following tamoxifen treatment (VCP^RC/FL^; Rosa26-Cre ERT2 (cVCP-RC) (Figure S5) (paper in print). Similar to our previous study, lysates from the cortex of VCP^RH/WT^ have no changes in the levels of autophagic (LC3, Sequestosome-1/p62), ER stress (BiP/GRP78), or ubiquitinated proteins (Figure 5A-B). Following five days i.p. tamoxifen treatment, cortical lysates from cVCP-RC mice have a 41% reduction in total VCP protein level but no changes in autophagic levels (LC3, Sequestosome-1/p62), ER stress (BIP/GRP78) or ubiquitinated proteins (Figure 5A-B). However, high molecular weight ubiquitinated proteins and SQSTM1 levels increase with age as demonstrated by immunoblot of cortical lysates at 6 months supporting that VCP dysfunction is present (Figure 5C-D). We have previously demonstrated that an increase in Gal3 levels occurs early before autophagic dysfunction in VCP^RH/WT^ mouse muscle ^17^. Similar to skeletal muscle, Gal3 and LAMP1 levels are increased in both VCP^RH/WT^ and cVCP-RC mouse cortical lysates suggesting an accumulation of damaged late endosomes (Figure 5A-B) ^15, 17^.

**Figure 5:**
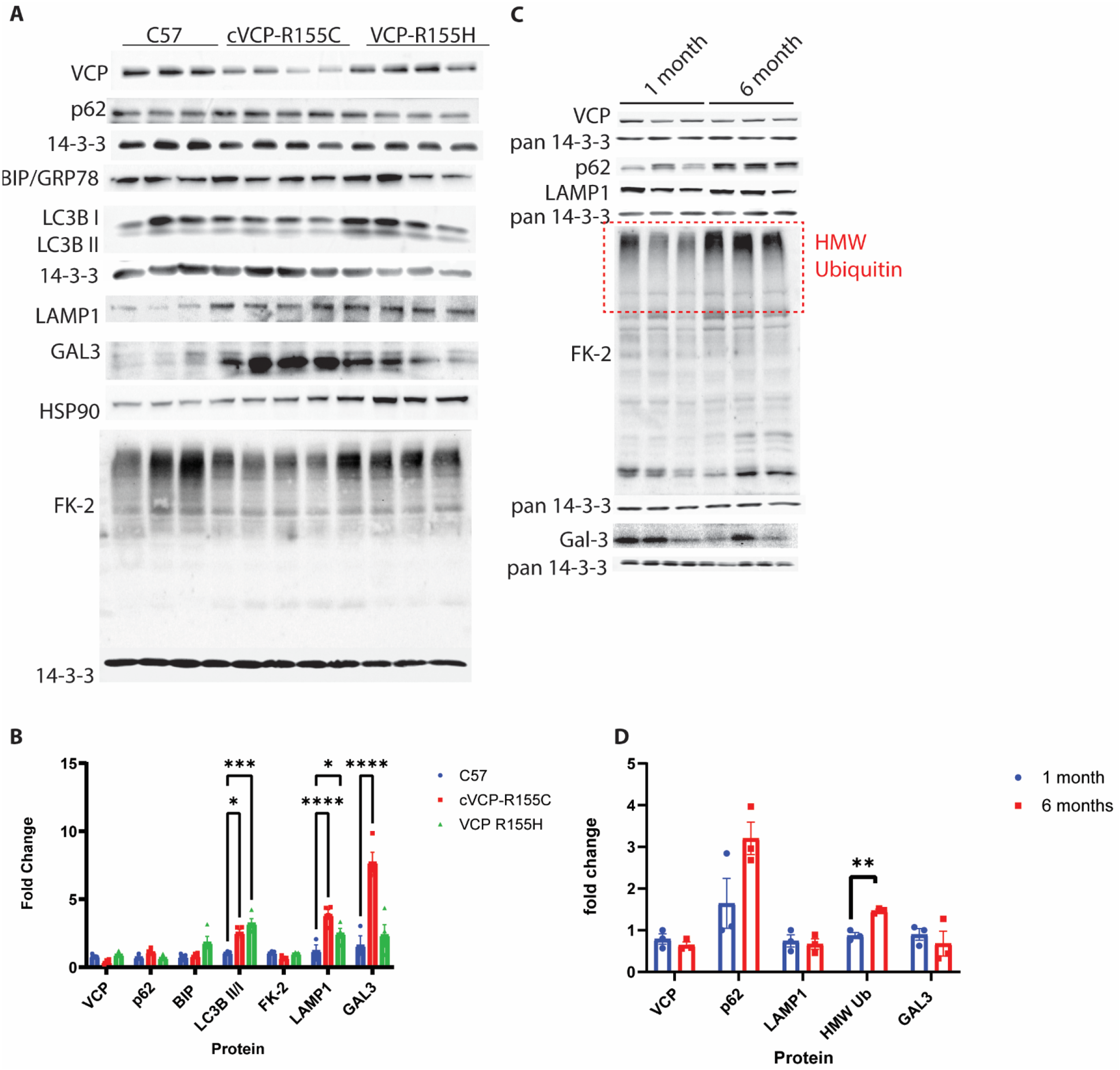
VCP disease mutant mice accumulate Galectin-3. (A) Representative immunoblot for VCP, p62, 14-3-3, BiP/GRP78, LC3, Lamp1, Galectin-3, HSP90, and ubiquitin (FK2) from cortical brain lysates of 4 month old C57 (n=3), VCP^R155H/WT^ (n=4) and cVCP-R155C (n=4) mice. In the case of cVCP-R155C mice, they are intraperitoneally injected with tamoxifen at 90 days of age and the brain was collected after one month. (B) Quantification of band intensities of VCP, p62, Bip/GRP78, LC3, FK2, Lamp1 and Gal3. (C) Representative immunoblot for VCP, p62, 14-3-3, Lamp1, Galectin-3, and ubiquitin (FK2) from cortical brain lysates of cVCP-R155C mice following one month or six months of i.p. tamoxifen treatment. (D) Quantitation of band intensities of VCP, p62, Lamp1, FK2 and Gal3 (n=3 for both groups).

We injected 5ug αS PFF or PBS into the striatum of 4-month-old C57 control, VCP^RH/WT^, or cVCP-RC mice and harvested the brain after 3 months (Figure 6A). Untreated control, VCP^RH/WT^, cVCP-RC mice, or mice treated with PBS had no phospho-αS staining in any brain regions (Figure 6A-E). In contrast, C57 control mice injected with αS PFF had a significant increase of phospho-αS in multiple brain regions (Figure 6A-E). This increase was significantly increased above that of injected C57 mice in the anterior and posterior cortices of VCP^RH/WT^, and cVCP-RC injected with αS PFF (Figure 6A-E). Other brain regions such as the amygdala and substantia nigra trended toward an increase in phospho-αS staining but did not reach statistical significance (Figure 6F-G).

**Figure 6:**
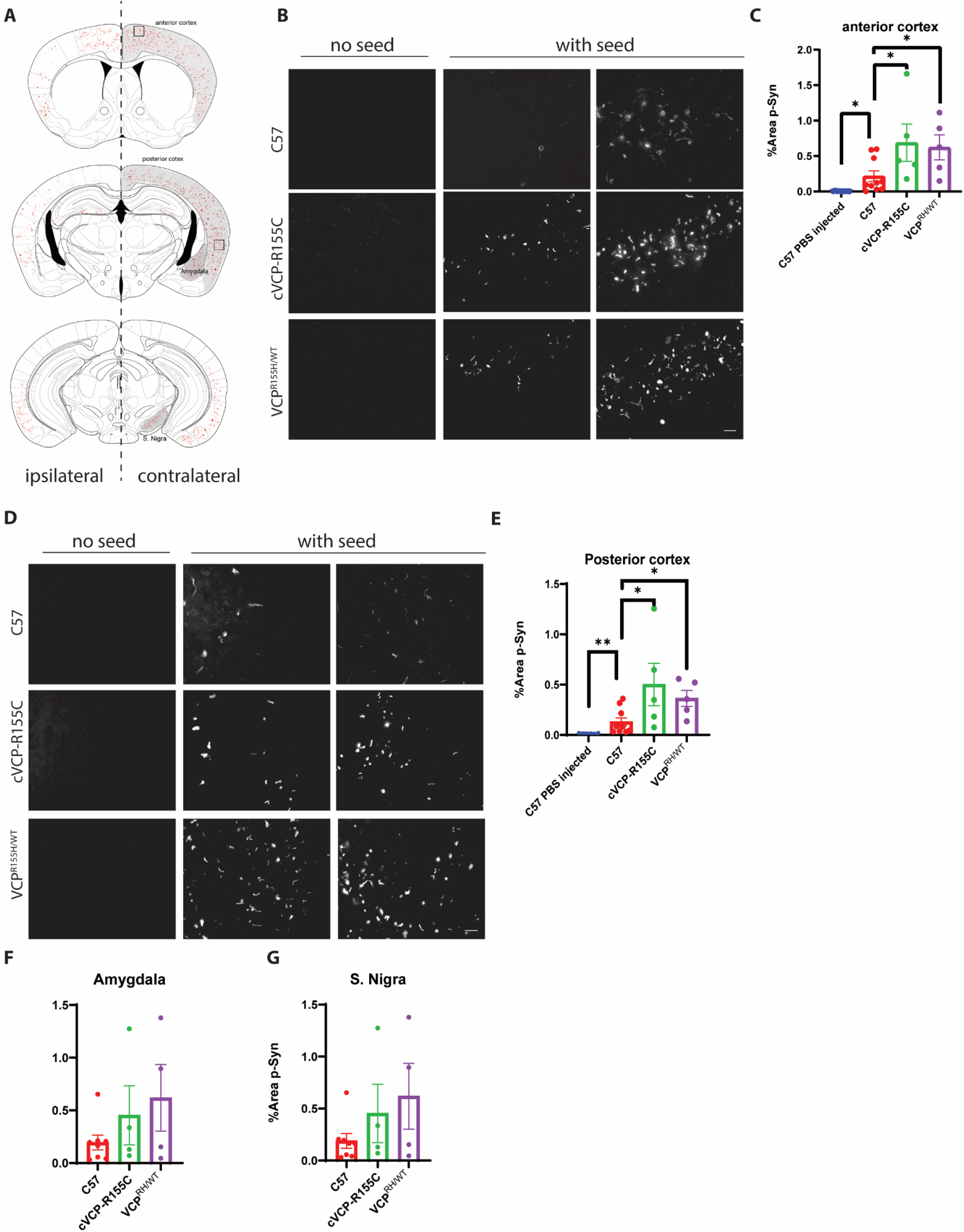
VCP disease mutations enhance αS seeding *in vivo*. (A) Schematic of coronal sections through mouse brains. Shaded regions indicate areas utilized for quantitation and boxes denote regions corresponding to images in B and D. Red dots represent p-syn detected. (B) Representative immunofluorescence images with pSer129-syn antibody of anterior cortices from C57, cVCP-R155C, and VCP^R155H/WT^ mice injected unilaterally into the striatum with 5ug αS PFF after 90 days. 4 month aged C57 (n=10), VCP R155H/WT (n=5) and VCP R155C/FL; Rosa26-Cre ERT2 (cVCP-R155C) (n=5). Scale bar=25 μm. (C) Quantitation of the percentage of p-syn in entire anterior cortices (* P<0.05 by student’s t-test). (D) Representative immunofluorescence images with pSer129-syn antibody of posterior cortices from C57, cVCP-R155C, and VCP^R155H/WT^ mice as described in A. scale bar=25 μm. (E) Quantitation of the percentage of p-syn in posterior cortices (* P<0.05, **P<0.01(* P<0.05 by student’s t-test). (F-G) Quantitation of p-syn in Amygdala and substantia Nigra. (ns, no significance student’s t-test)

### VCP disease mutations enhance TDP-43 seeding

A subset of VCP patients have Parkinsonism and post-mortem evidence of αS pathology, however, most patients have TDP-43 inclusions in the CNS and muscle ^8–10^. To evaluate the role of VCP in the seeding of TDP-43, we developed a TDP-43 seeding assay in primary hippocampal neurons. The addition of TDP PFF to HNs resulted in the appearance of phosphorylated TDP-43 Ser409/410 (pTDP) positive puncta in a concentration-dependent manner (Figure 7A-B) and redistribution of pTDP from the nucleus to the cytoplasm (Figure 7C) after 5 days. pTDP staining was not increased or altered when HNs were treated with αS PFFs (Figure S6). Fractionation of lysates from HNs one or 5 day post-treatment with buffer, non-aggregated TDP-43 (monomeric TDP-43), or TDP-43 aggregated at room temperature for 24 hours (TDP-43 PFF) and subsequent immunoblot for pTDP revealed an increase in high molecular weight TDP-43 in the RIPA insoluble fraction of TDP PFF treated HNs (Figure 7D). In addition, TDP-43 PFF induced cytosolic pTDP-43 puncta that co-localized with Sequestosome-1 and TIA1 similar to pathologic TDP-43 inclusions in patients (Figure 7E). As seen with VCP mutant expression in TDP-43 biosensors, treatment of primary HNs from VCP^WT/WT^ and VCP^RH/WT^ embryos with TDP-43 PFF revealed an increase pTDP-43 puncta in VCP^RH/WT^ HNs compared with VCP^WT/WT^ HNs (Figure 7F-G).

**Figure 7:**
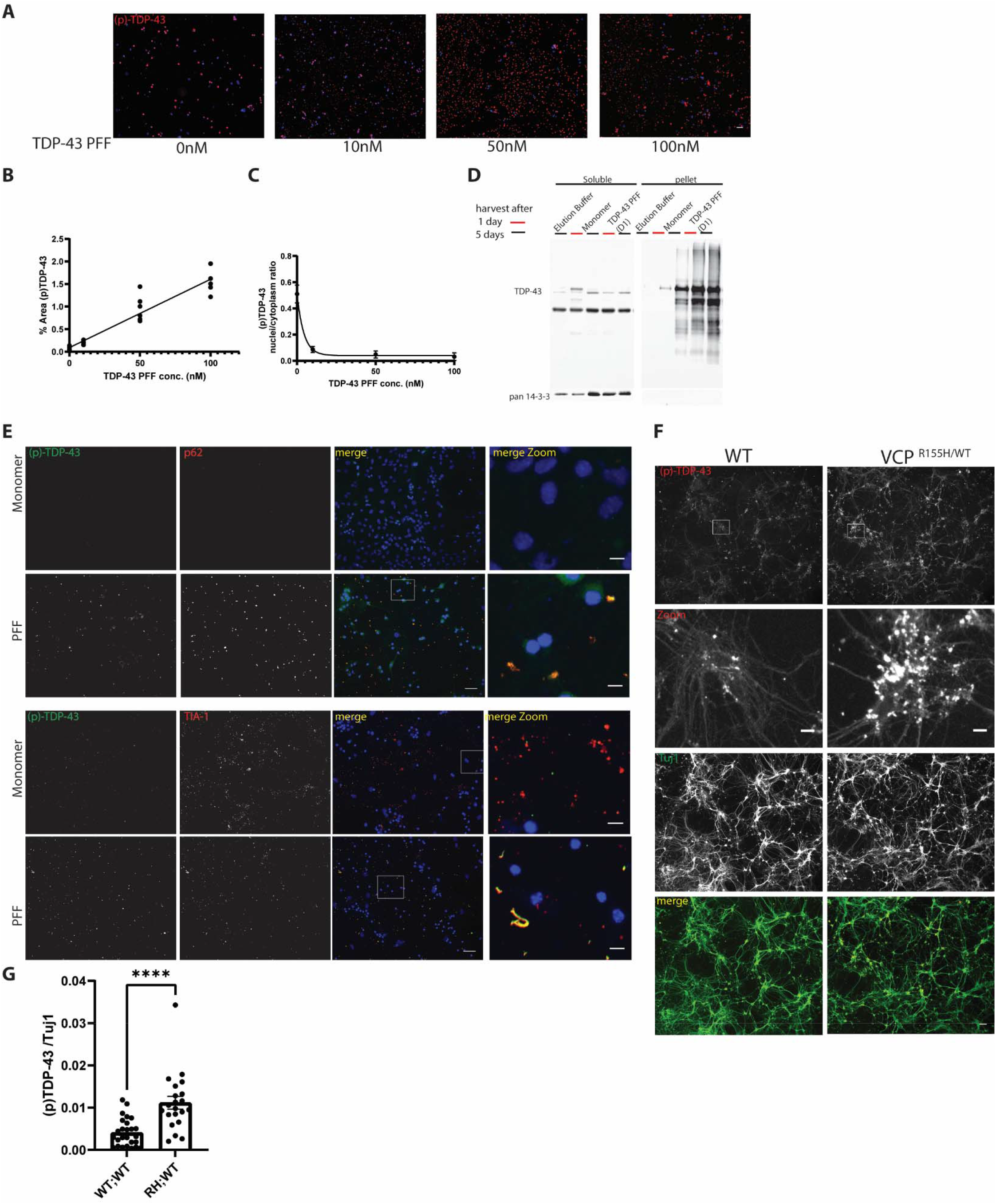
VCP inhibition or VCP disease mutations enhance TDP-43 seeding in cells and neurons. (A) Immunofluorescent staining for pTDP-43 (red) and nuclei (blue) in primary hippocampal neurons treated with varied concentrations of TDP-43 PFF for 5 days. Scare bar= 50μm (B) Quantitation of area of pTDP-43 immunofluorescence as in 7D. n ≥5 for each concentrations. (C) Quantitation of the ratio of nuclear to cytoplasmic fluorescence of pTDP-43 as in 7D. pTDP-43 fluorescence intensity was measured separately in nucleus and cytoplasm. More than 45 nucleus are counted in each group. (D) Immunoblot for TDP-43 from detergent soluble and insoluble lysates of HNs treated with TDP-43 monomer or PFF and then harvested after 1 or 5 days. Note that the 2%SDS insoluble fraction has high molecular weight TDP-43 positive multimers. 14-3-3 is a loading control. (E) Immunofluorescent images for pTDP-43 (green), SQSTM1 (red upper panels), TIA1 (red lower panels) and DAPI nuclei (blue) from neurons treated with TDP-43 monomer or PFF for five days. Scare bar= 50 and 10 μm for original and zoom-in images. (F) Immunofluorescence for phospho-TDP-43 (red) and Tuj1 (green) in HNs form wild-type mice or mice carrying a VCP-R155H knockin allele (VCPR155H/WT) treated with TDP-43 for PFF for 5 days. (Scare bar= 50 and 10 μm for original and zoom-in images). (G) Quantitation of phospho-TDP/Tuj1 staining as in 7I (Neurons coming from 3 and 4 independent cultures from WT and VCP^R155H/WT^ embryos. Outlier is removed by ROUT method, Q=1%, followed by Student’s t test. n=26, 21 for WT and VCP^R155H/WT^ group respectively. **** P<0.0001.)

## DISCUSSION

Functional genomic screens utilizing CRISPR knockout approaches are an invaluable resource to elucidate proteins related to distinct cellular pathways. Here, we employed a CRISPR whole-genome KO screen to identify modifiers of αS seeding using an αS biosensor cell line. αS seeding spans many cellular processes that include endocytic uptake, vesicular trafficking, templated aggregate conversion, and protein degradation by both the proteasome and autophagy. Our screen identified proteins associated with vesicular trafficking between the ER, golgi and endosome, and the cellular stress response. Notably these pathways have been identified as modifiers of αS toxicity and stability in yeast and cell models ^31, 32^. Vesicular trafficking may have been particularly enriched since our screen and the αS biosensor cell line required that αS PFFs be applied with the carrier, Lipofectamine, as a means of facilitating endocytic uptake.

Our further studies expanded upon the role of VCP in αS seeding. VCP is a multifunctional protein necessary for many ubiquitin dependent processes that include protein degradation, vesicle trafficking, cell division and organelle clearance ^7^. Recently, we identified a role for VCP in the recognition of permeabilized late endosomes and their subsequent lysophagic degradation ^15^. Endocytosed proteopathic seeds such as αS and Tau enter the cytoplasm, where templated aggregate conversion of soluble monomer occurs, by damaging the endosomal membrane ^5, 15, 33^. The fate of permeabilized late endosomes depends upon the degree of membrane damage. For example, some damaged endosomes are rapidly repaired by ESCRT proteins ^34^. In contrast, endosomes damaged beyond repair are tagged by intracellular galectins such as galectin-3 ^15^. Galectin positive endosomes recruit the ubiquitin ligase, Trim16 which ubiquitinates endosomal membrane proteins ^35^. Notably, only lysine-63 linked ubiquitin chains on the endosome surface are targeted for lysophagy ^15^. VCP in association with UBXD1, PLAA and the deubiquitinase YOD1 recognize and cleave lysine-48 linked ubiquitin chains on damaged late endosomes leaving lysine-63 linked ubiquitin chains allowing lysophagic degradation ^15^. Loss of VCP or VCP disease mutant expression leads to the persistence of galectin-3 positive damaged late endosomes in cells, mouse models and patient tissue ^15, 17^.

A previous genomic screen using a Tau biosensor line identified several components of the ESCRT machinery as suppressors of Tau seeding ^36^. Our screen identified VCP and Trim16 as suppressors of αS seeding. Further experiments found that knockdown of the VCP adaptor, UBXD1 that is necessary for lysophagy and VCP disease mutations defective in lysophagy also increase αS seeding. One distinction between these two screens is the use of lipofectamine to facilitate entry into the endocytic pathway of αS. Lipofectamine is known to damage endosomal membranes and may allow αS to generate larger “holes” that are not repaired by ESCRTS ^34^. However, VCP’s role in seeding was not exclusively lipofectamine dependent since VCP inhibition, knockdown or VCP mutant expression facilitated “naked” αS PFF seeding in HNs and *in vivo*.

VCP disease mutations cause multisystem proteinopathy (MSP) ^37^. MSP is a late onset degenerative disorder with varied phenotypes and pathologies. These include inclusion body myopathy, ALS and FTD ^18^. While the predominant aggregate pathology in the brain is reported to be TDP-43, several studies support the identification of αS positive aggregates in the brain ^8–10^. Indeed, ~5% of MSP patients have coincident Parkinsonism ^19^. Notably, two recently identified families with a VCP-D395G mutation were found to have distinctive tau pathology leading to the description of a vacuolar tauopathy in the CNS ^11^. Aggregate pathology in the skeletal muscle can be varied and include TDP-43, hnRNPA1/A2B1, SQSTM1, β-amyloid, desmin and VCP ^38–40^. Weakness typically precedes the onset of neurodegenerative features such as dementia by 10 years suggesting that pathology begins in peripheral tissue such as skeletal muscle ^19^. Whether protein aggregates from skeletal can seed the aggregate process in motor neurons or cortical neurons remain speculative. It is noteworthy that mice carrying the D395G missense mutation in VCP had an increase in Tau seeding supporting that VCP disease mutations can facilitate the propagation of different aggregate species ^11^.

TDP-43 inclusions are a prominent feature in affected VCP disease tissue ^9, 38^. Our data further supports that TDP-43 PFF similar to αS-PFF can seed pathologic TDP-43 inclusions in HNs. TDP-43 seeding in HNs recapitulates several features of TDP-43 pathology such as phosphorylation, cytoplasmic redistribution and co-localization with stress granule markers and SQSTM1 ^41^. VCP disease mutation expression increases pathologic TDP-43 inclusions as measured by an increase in phospho-TDP-43 immunostaining in HNs. This process is TDP-43 PFF dependent since monomeric TDP-43 fails to have the same effect. Whether the accumulation of TDP-43 inclusions is also affected by an additional role for VCP in stress granule clearance remains to be determined ^42^.

## METHOD AND MATERIAL

### αS FRET seeding assay

Generally, HEK 293T αS-CFP/YFP is plated in a black-bottomed 96-well plate with the density of 80k/well in DMEM media with 10% FBS and Penicillin-Streptomycin. Three control cell lines – no-transfected HEK293T cells, αS-CFP, and αS-YFP transduced cells are cultured in the same condition. αS PFF is sonicated and prepared with OPTIMEN and 1ul Lipofectamine 20000 (Invitrogen) for each well and add dropwise to the cell after 48 hours. The cells are harvested after 24 hours for flow cytometry, same as reported. Briefly, the cells are detached by 0.05% trypsin-EDTA (Gibco), centrifuged, then fixed with 2% PFA for 15 minutes, and finally re-suspended in MACSima Running Buffer. The samples are analyzed by MACSQuant® VYB. FRET signal is excited by 405nm lasers and detected by 525/50 band pass filter. While the CFP and YFP are excited by 405nm and 488nm lasers and filtered by 450/50nm and 525/50nm, respectively. The data is analyzed with FlowJ v10 software. Each FRET signal is calculated as percentage of FRET-positive cell timing Median FRET fluorescence intensity, and then normalized to its vehicle control.

VCP mcherry vectors are a gift from Hemmo Meyer’s lab. The mutations is confirmed by Sanger sequencing (GENEWIZ) with VCP plasmid primers described before. 250ng of the plasmid is transfected with OPTIMEN and 0.5ul Fugene 6 (Promega) in each well 24 hours after plating. The cells then treated the same way as described above. The mcherry signal is excited by 561nm laser and filtered via 615/20nm. The FRET signal is analyzed separately for mcherry positive and negative cells.

Knockdown is achieved by reverse transfection of Human SMARTPOOL siRNA from Dharmacon or Thermo Silencer Select siRNA. 6pmol siRNA is prepared in OPTIMEN and 0.3ul Lipofectamine™ RNAiMAX (Invitrogen) according to its protocol and add to each well in a 96-well plate. Then 80k suspended αS-CFP/YFP cell is plated in each well already with siRNA droplet. The αS PFF is treated 48 hours after plating as described above.

### Genome-wide CRISPR-Cas9 screens on αS biosensor line

αS-CFP/YFP HEK293T cells were first transduced with WT cas9-blast.Single clone were sorted and cultured. The new cas9 αS CFP/YFP line maintained the both αS-CFP and αS-YFP, and capable of seeding. Cas9 function were validated by a synethic gRNA and obtained with 99% NHEJ activity. About 50 million HEK293T syn CFP/YFP Cas9-blast cell were plated and then infected with pooled lentivirus with Burnello gRNA library (Addgene #73178-LV) with 8ug/ml polybrene (MOI=0.3) the next day. After 24 hours, cells would undergo 1ug/ml puromycin selection. Selected cells were replated at density of 6.4*10^5 cell/ml after 96 hours of selection and replaced with fresh puromycin. 2 days later, harvest 1/5 of the cell (~20 million) (untreated group, for library representation) and seeded the rest with 10 nM αS -PFF. The seeded cells were collected the same as normal αS FRET assay as described above after 24 hours and sorted by Sony SY3200 cell sorter. DNA extraction were performed via QIAamp DNA Blood Midi on FRET positive and negative cells, as well as unsorted cells were separately,amplified by PCR, and deep sequenced by Illumina NovaSeq. The FRET positive and negative groups were compared with untreated total population group separately via Megack RRA. For pathway enrichment, 153 hits were input to g:profiler and plot via cytoscape as described before.

### αS and TDP-43 Fibril preparation

αS PFF and monomer is generated as described before ^21^. Briefly, purified recombinant αS monomer (2 mg/ml) was incubated in 20 mM Tris-HCl, pH 8.0, 100 mM NaCl for 72 h at 37°C with shaking at 1000 rpm in an Eppendorf Thermomixer. To determine the concentration of fibrils, the fibril reaction mix was centrifuged at 18,000×g for 15 min to separate fibrils from monomer. The concentration of αS monomer in the supernatant was determined in a BCA protein assay according to the manufacturer’s instructions, using a bovine serum albumin (BSA) standard curve. The measured decrease in αS monomer concentration was used to determine the concentration of fibrils in the 72 h fibril reaction mixture. To isolate pre-formed fibrils (PFF) from monomer, centrifuge the αS mix at 18,000×g for 15 min to separate fibrils from monomer. Resuspend fibril pellet in the buffer containing 20 mM Tris-HCl, 100 mM NaCl, pH 8.0. αS PFF always freshly sonicated right before seeding.

Fluorescently labeled fibrils of αS were generated as previously described ^43^. αS (1 mg/mL) was dissolved in 100 mM NaHCO3, sonicated for 15 min, and spun through a 50 kD filter (Amicon UFC5050) at 16,100 ×g for 15min. Alexa Fluor 647 NHS Ester (Thermo Fisher A20006) was dissolved in DMSO to 10 mg/ml. Dye solution (molar ratio of dye: αS = 2.1:1) was pipetted into monomerized αS during stirring and the mixture was stirred on bench for ~1h. The mixture was then loaded onto a size exclusion column (Superdex 75 10/300 GL) and eluted with 5mM NaOH. The peak containing monomeric, labelled αS was collected, aliquoted and kept frozen until use. For aggregation assays, αS was dissolved in 10 mM NaOH at 1 mg/mL. αS-647 was added at 5% labelling ratio. Then the solution was sonicated for 20 minutes, filtered through a 100 kD membrane filter (Amicon Ultra, 540655) at 16,100 × g for 15 min at 4°C. The protein and dye concentrations were measured by absorption at 280 nm and 647 nm, respectively, and the labelling ratio was determined to be 4.9%. To prepare labelled αS fibrils, monomer solution (5% α-syn-AF647) and solutions were incubated in 100 mM NaP, pH 7.4, 10 mM NaCl for 120 h aggregated in a non-binding 96 well plate (Corning, #3651) at a concentration of 30 μM with intermittent shaking in aggregation buffer (100 mM NaP, pH 7.4, 10 mM NaCl). A 2 mm diameter glass bead was added to each well to accelerate the aggregation through stirring. The plate was kept at 37°C and agitated by orbital shaking once every 1 minute for 5 seconds.

Recombinant TDP-43 (rTDP-43) was generated in *Escherichia coli* and purified as previously described. Briefly, rTDP-43 was bound to nickel–nitrilotriacetic acid–agarose and washed with wash buffer 1 (50 mM Tris, pH 8.0, 500 mM NaCl, 10% glycerol, 10% sucrose, 1 mM TCEP), washed with wash buffer 2 (50 mM Tris, pH 8.0, 500 mM NaCl, 10% glycerol, 10% sucrose, 50 mM Ultrol Grade imidazole, pH 8.0, 1 mM TCEP), and finally eluted (50 mM Tris, pH 8.0, 500 mM NaCl, 10% glycerol, 10% sucrose, 300 mM Ultrol Grade imidazole, pH 8.0, 1 mM TCEP). Then, rTDP-43 was ultracentrifuged in a Beckman Coulter Optima MAX-XP Ultracentrifuge at 40,000 rpm for 30 min at 4 °C to remove any pre-existing aggregates. Soluble protein was diluted to 4uM in the reaction buffer (50 mM Tris, pH 8.0, 250 mM NaCl, 5% glycerol, 5% sucrose, 150 mM Ultrol Grade imidazole, pH 8.0, 0.5mM TCEP). rTDP-43 aggregation was started by shaking at 1,000 rpm at 22 °C for 30 min with an Eppendorf ThermoMixer C. Samples were incubated at 22 °C and collected after one to ten days. Full length TDP-43 recombinant protein is produced as previously described ^21^. To obtain TDP-43 monomer, TDP-43 protein was ultracentrifuge 40,000g 30 mins at 4 °C. The supernatant was collected and freshly used.

### Primary neuronal culture

WT hippocampal neurons were obtained from E17-18 mice (Charles River). Hippocampi are dissected in calcium- and magnesium-free Hanks’ Balanced Salt solution (HBSS) and dissociated by 0.05% Trypsin-EDTA at 37C for 5-10 mins and followed by 1% DNase I for 2 mins ^44^. The cells are then resuspended with plating media to the concentration of 125k/ml and plated on Poly-D-lysine coated plates or coverslips. The media is changed to neurobasal media (neurobasal plus + B27 + 5mM L-Glutamine) after 2-4 hours. 1mM Ara-C is added to inhibit the growth of glia. The αS PFF is sonicated and add directly to the cell at DIV10.

VCP and UBXD1 shRNA is delivered via lentivirus. Lentivirus were added to the neurons at DIV5 with MOI≥5. For LLoMe or VCP inhibitors experiments, drug or vehicle control was added at the same time with αS PFF for 4 hours, and then the media is fully exchanged to the conditioned media without drug and αS. The neurons were harvested at DIV15.

R155H/WT neurons were cultured from embryos from R155H/WT intercross. The hippocampus from each embryo was dissected and cultured separately and then plate at the same density.

The genotypes were examined by PCR (Transnetyx) (Forward Primer: CCTCTAATTGCACTTGTATTGCTTTGT; Reverse Primer: CTGGGATCTGTCTCTACAACTTTGA).

### Immunohistochemistry

Cells were fixed in 4% PFA for 10 mins and permeabilized with 0.1% Triton X-100 in PBS for 10 mins. Then the cells were blocked with 2% BSA in PBS at RT for 1 hour. Cells were stained with primary antibody at 4 °C overnight, followed by 3 washes with PBS. Cells were then incubated with the Alexa 488 555 or 647 tagged secondary antibody in 1:500 dilution for 1 hour at RT. The nucleus was stained with DAPI (1:1000) for 10 min at RT. After 3 wash with PBS, the cells are mounted by Mowiol. Pictures were taken by either Nikon Eclipse 80i fluorescence microscope or a Hamamatsu NanoZoomer. The images were process via ImageJ or NDP.view2. The antibody is listed in Appendix.

### Immunoblot

Mouse cortex or cells are lysed in RIPA buffer with protease inhibitor cocktails (PMSF and PIC) followed by two 30 sec on and 30 off sonication cycle at 50% power. The protein concentration is normalized by the BCA assay. Samples were loaded into 10 to 15% gel and transferred into nitrocellulose or PVDF membrane. The membranes were blocked by 5% milk in PBS-0.2% Tween20 and incubated with primary antibody in blocking solution overnight at 4 °C degree. Membrane then were washed three time with PBS-0.2% Tween20, and incubated with secondary goat anti-rabbit, mouse HRP antibody (1:5000) for 1 hour. Blot were rinsed three time with PBS-0.2% Tween20 and probed by fresh mixture of ECL reagents at dark and then exposed by SYNGENE.

To fractionate insoluble portion of αS, we preformed sequential extraction as described ^45^. Briefly, neurons were first dissolved in TBS-1%Tx-100, and sonicated for 10 cycles of 30s on, 30s off with 50% power. The lysate would incubate on ice for 30 mins. 1/10 of the lysate were saved as total protein, while the remaint was ultracentrifuged 100,000g 4 °C for 30 mins. The supernant was collected as Tx-100 soluble fraction.the pellet was wash with TBS-1%Tx-100, sonicated and ultracnetrifuged. Ultimately, the pellet was resupsended with TBS-2%SDS, and sonicated for 15 cycles of 30 s on, 30 s off. Soluble and insoluble fraction were run by western blot as normal. The loading amount were determined by the concentration of total protein measured by BCA.

TPD-43 soluble and insoluble extraction is done as our previous method. Briefly, neurons from one 6 well were first lysed with RIPA buffer with protease inhibitor cocktails (RIPA buffer) on ice. The lysate then sonicated with QSONICA sonicator for 10 cycles of 30 s on, 30 s off with 50% power. 1/10 of the lysate were saved as total protein. The rest was ultracentrifuged at 100,000g 4 °C for 30 mins. The supernatant was kept as RIPA soluble fraction. The pellet was then wash with RIPA buffer once, resonicated and ultracentrifuged with the same condition. The pellet finally resupsend with same amount of UREA buffer (30 mM Tris, pH 8.8, 7M urea, 2M thiourea, and 4% CHAPS) as insoluble fraction. Soluble and insoluble fraction were run by western blot as normal. The loading amount were determined by the concentration of total protein measured by BCA.

### Intrastriatal injection and mouse brain harvest

Both mice are Intraperitoneal injected with 75 mg tamoxifen/kg body weight at two months of age and wait one month for gene knockdown. The existence of the R155C mutation allele and VCP flop knockdown confirmed by PCR. The primers are listed in Appendix. Intrastriatial injection is performed as described. αS PFF is prepared as described above and diluted in sterile PBS. αS PFF is sonicated 10 mins before injection. The mouse is anesthetized and injected at the dorsal striatum (Bregma=0.2mm, midline=2.0mm, depth=−3.2mm) of the left hemisphere. The same amount of PBS is used as vehicle control. The recovery of mice is monitored in the following week and sacrificed after 90 days. Mouse was first anesthetized in Isofluorane chamber and perfused with PBS containing herapin. The whole brains were removed from the skull and fixed in 4% PFA overnight at 4 °C degree and cut coronally into 40micrometer sections for immunostaining.

### Statistical Analysis

The data (except CRISPR screening) is analyzed by GraphPad Prism 9. Statistical tests included unpaired t test, one-way ANOVA, multiple t-test, simple linear regression, nonlinear regression and two-way ANOVA. Data were displayed as mean ± SEM. Two-stage step-up method of benjamini krieger and yekutieli, Dunnett, sidak correction were used to minimize false alarm from multiple comparsion.

---

**Figure S1**: <u>spCas9-gRNA αS biosensor maintain normal seeding capacity.</u> spCas9 αS biosensor was treated with pooled library and selected as described in Fig 1A, followed by αS seeding at different concentration. After 24hours, cells are harvested and the percentage of FRET is measured by flow cytometry.

**Figure S2:** <u>αS biosensor shows seeding activities in a concentration dependent manner.</u> (A) αS biosensor was treated with αS PFF at different concentration for 4 or 24 hours. Cells are harvested after 24 hours and analyzed the same as 2A. (B) Alexa647 tagged αS PFF show moderate seeding capacity according to FRET assay.

**Figure S3:** <u>Monomer is insufficient to induce seeding under drug treatment or VCP KD.</u> (A) Representative pictures of HNs treated with αS monomer for 4 hours together with either LLoMe (1uM), ML240 or vehicle cotnrols at DIV10. The cells are harvested after 5 days and stained with p-syn and Tuj1. αS PFF treated HNs are presented as positive control. (B) Representative pictures of HNs of VCP KDs or scramble control treated with αS monomer at DIV10. The cells are harvested after 5 days and stained with p-syn and Tuj1. αS PFF treated HNs are presented as positive control.

**Figure S4:** <u>Gene knockdown efficiency and VCP vector transfection.</u> (A) αS biosensor is reverse transfected with siRNA (6pmol). Cell lysis are saved at 48 and 72 hours after transfection. Western blot is preformed on each KD for protein level change. (B) αS biosensor cells with different KD are transduced with Alexa-647 tagged αS PFF for 4 hours. The percentage of uptake is calculated the same way as 2F. (C) αS biosensor VCP mutated vectors showed similar level of VCP exogenous expression, tested by Western blot.

**Figure S5:** <u>Genotyping of cVCP-R155C mice.</u> Cortical tissues from 4 month old cVCP-R155C and C57 are genotyped. cVCP-R155C mice are i.p. injected with Tamoxifen in 5 continuous day and harvested after one month for genotyping.

**Figure S6:** <u>pTDP-43 is TDP-43 PFF specific in hippocampal neurons.</u> TDP4-43 monomer, TDP-43 PFF or αS PFF is added as indicated. After 5 days, HNs are harvested and stained with (p)TDP-43 and (p)syn(81A)

## Notes

### Competing Interest Statement

The authors have declared no competing interest.

