## supplement figure 1-6 for "VCP protects neurons from proteopathic seeding"

### Slide 1
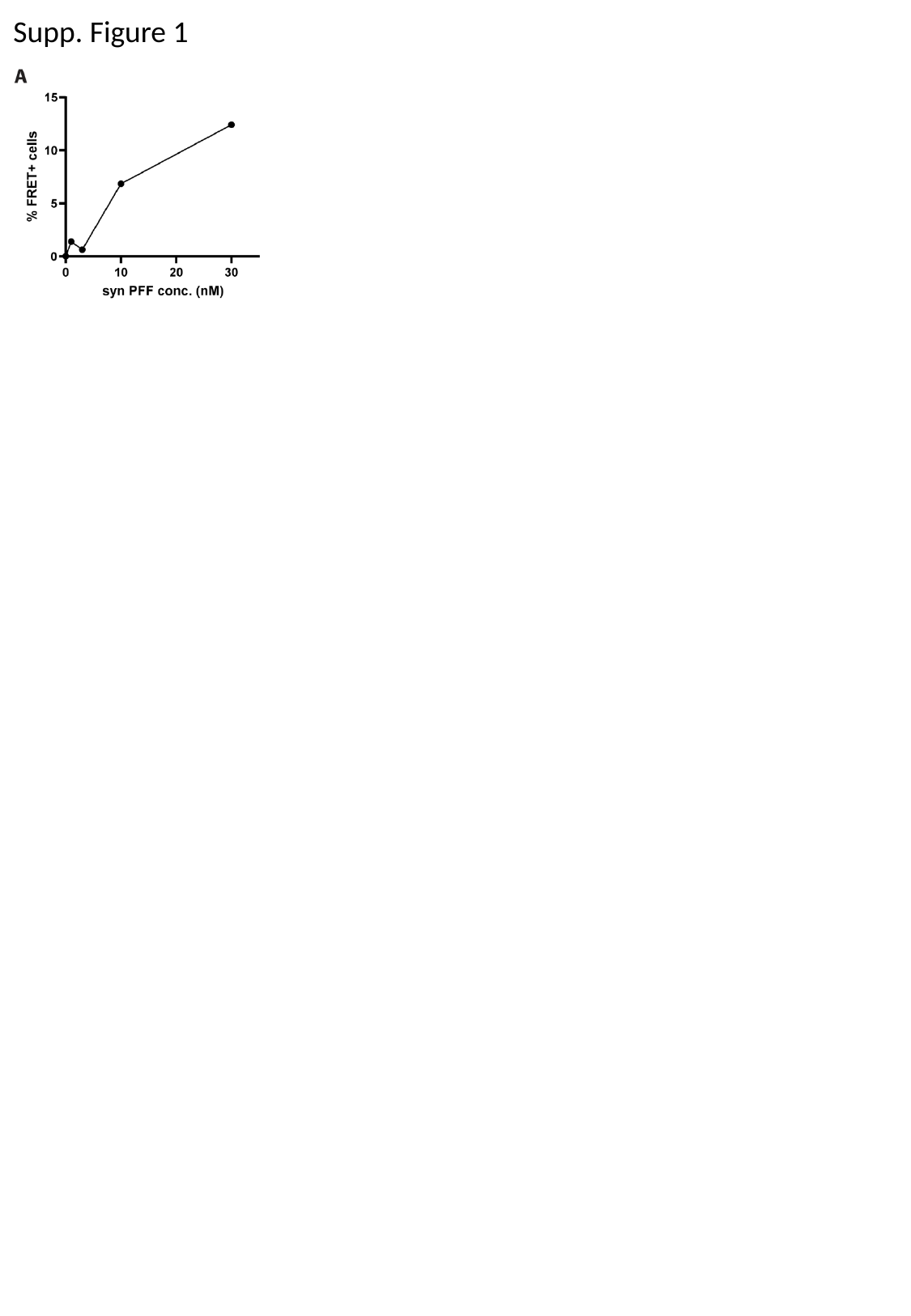

Supp. Figure 1

### Slide 2
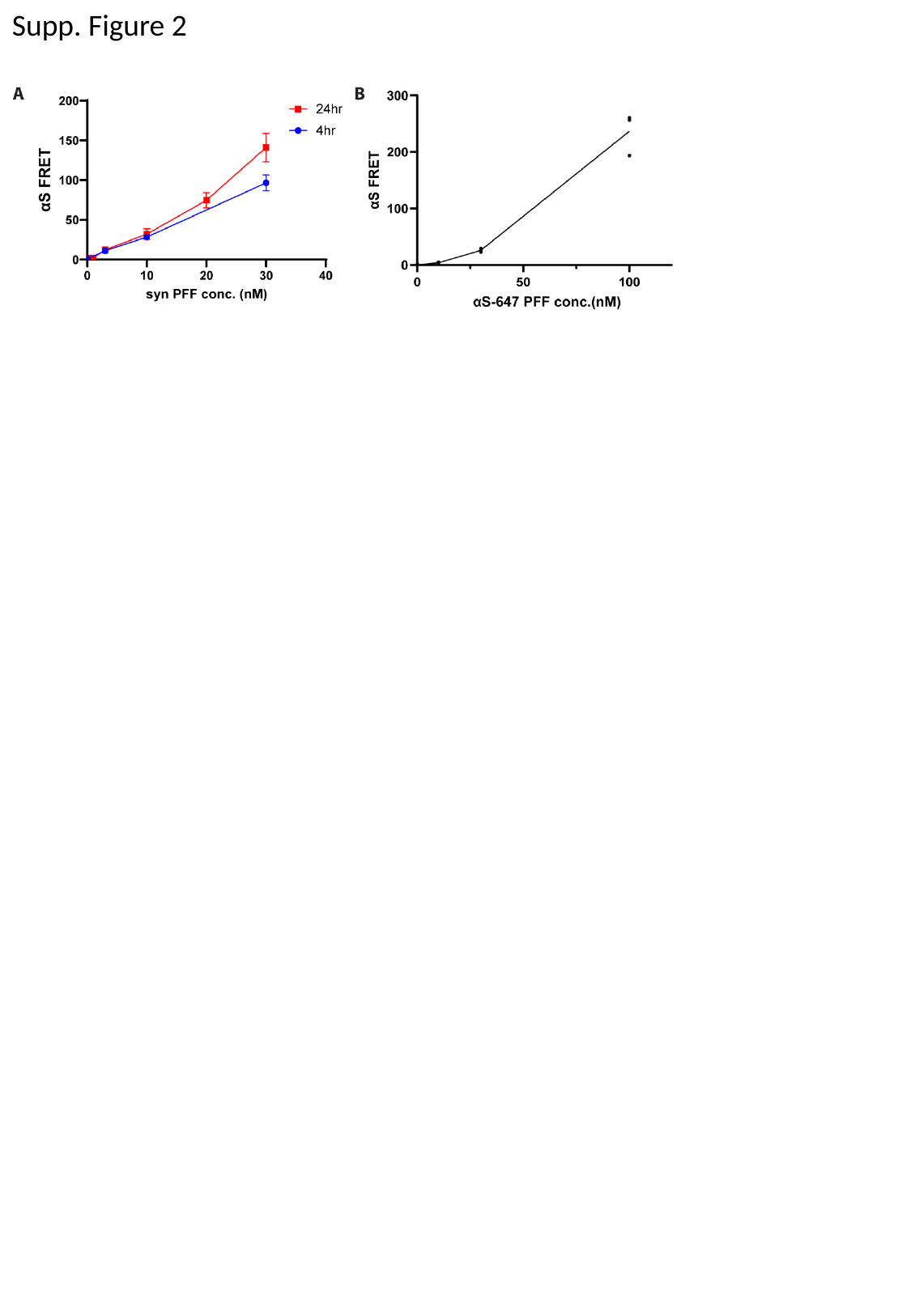

Supp. Figure 2

### Slide 3
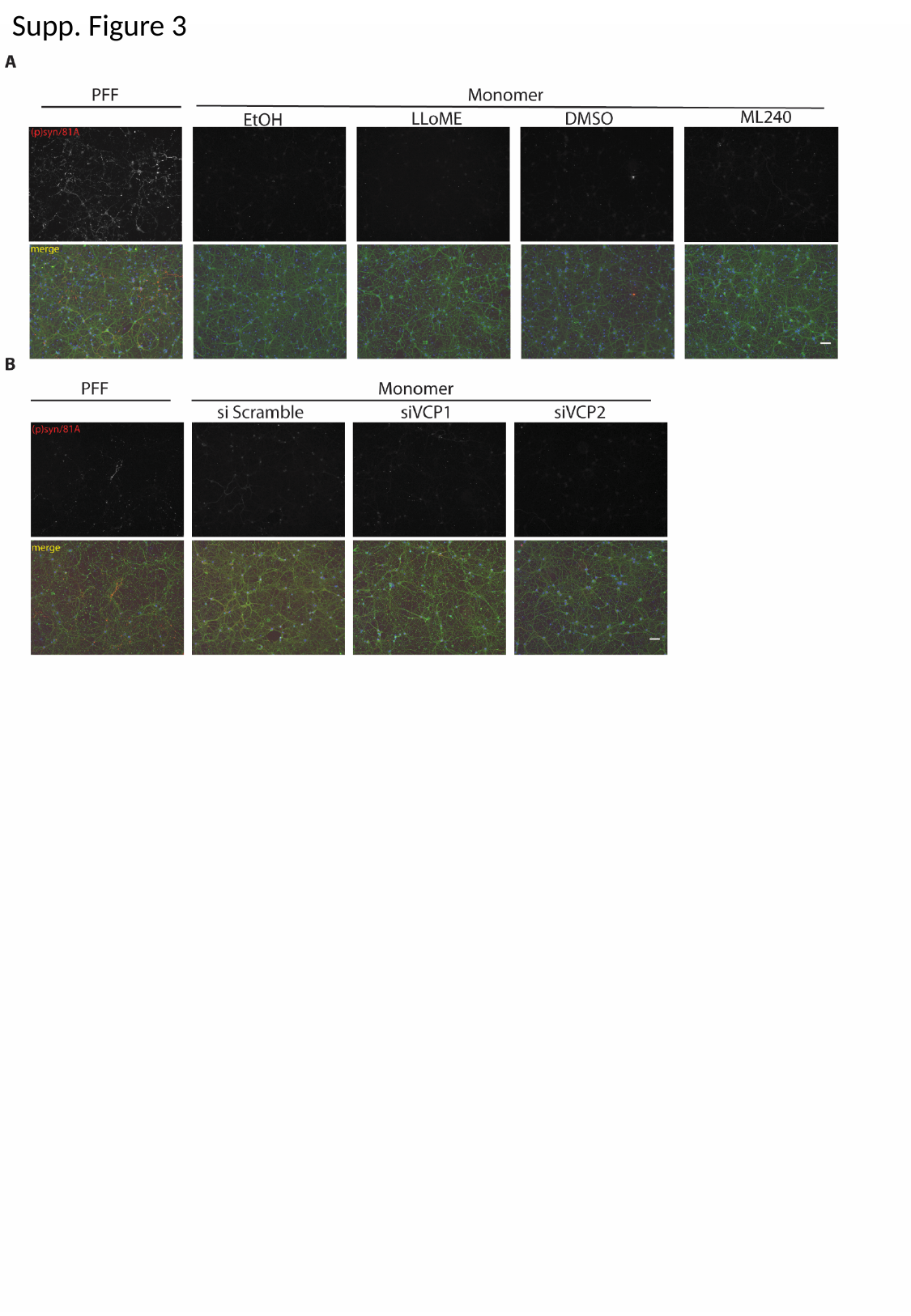

Supp. Figure 3

### Slide 4
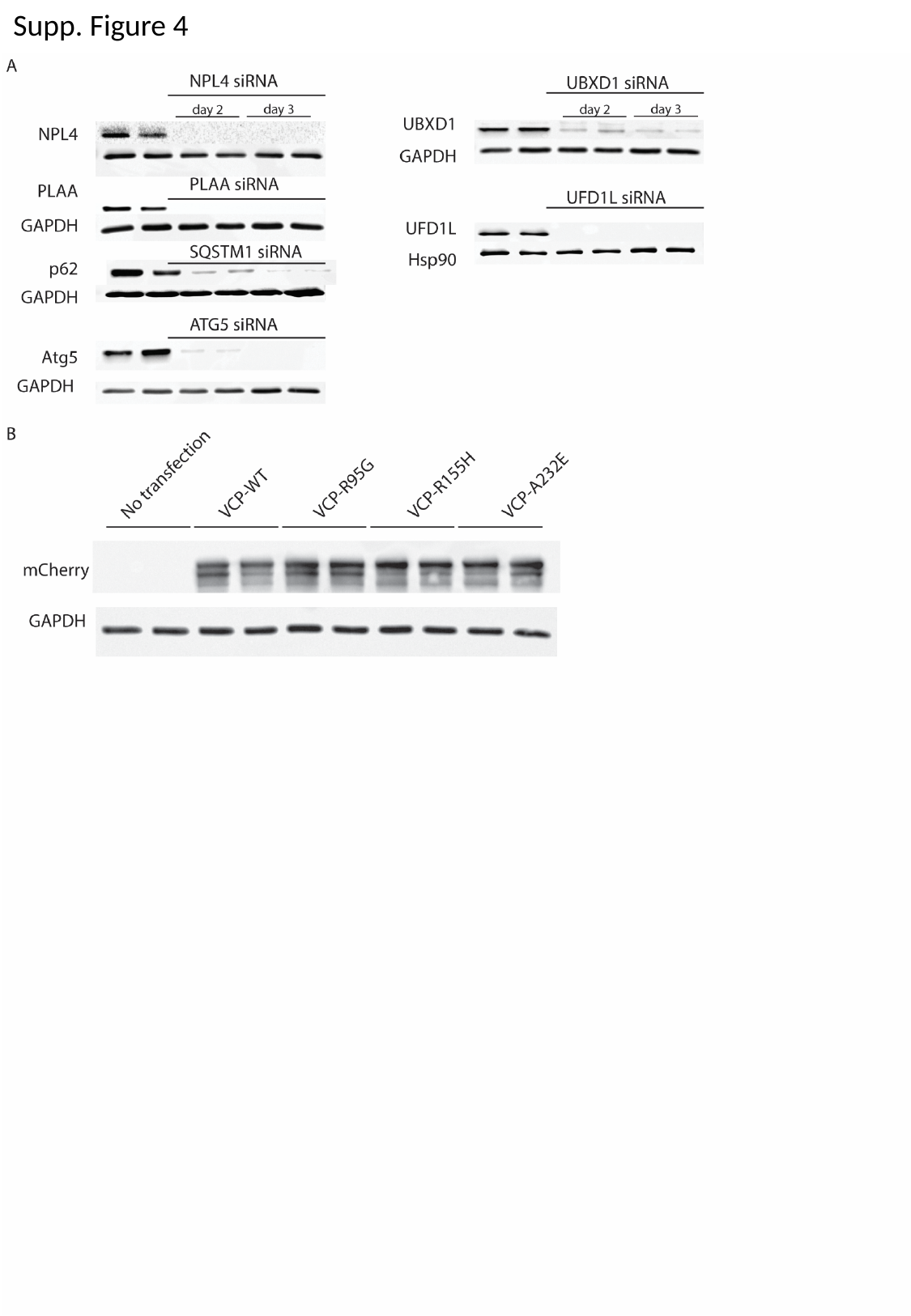

Supp. Figure 4

### Slide 5
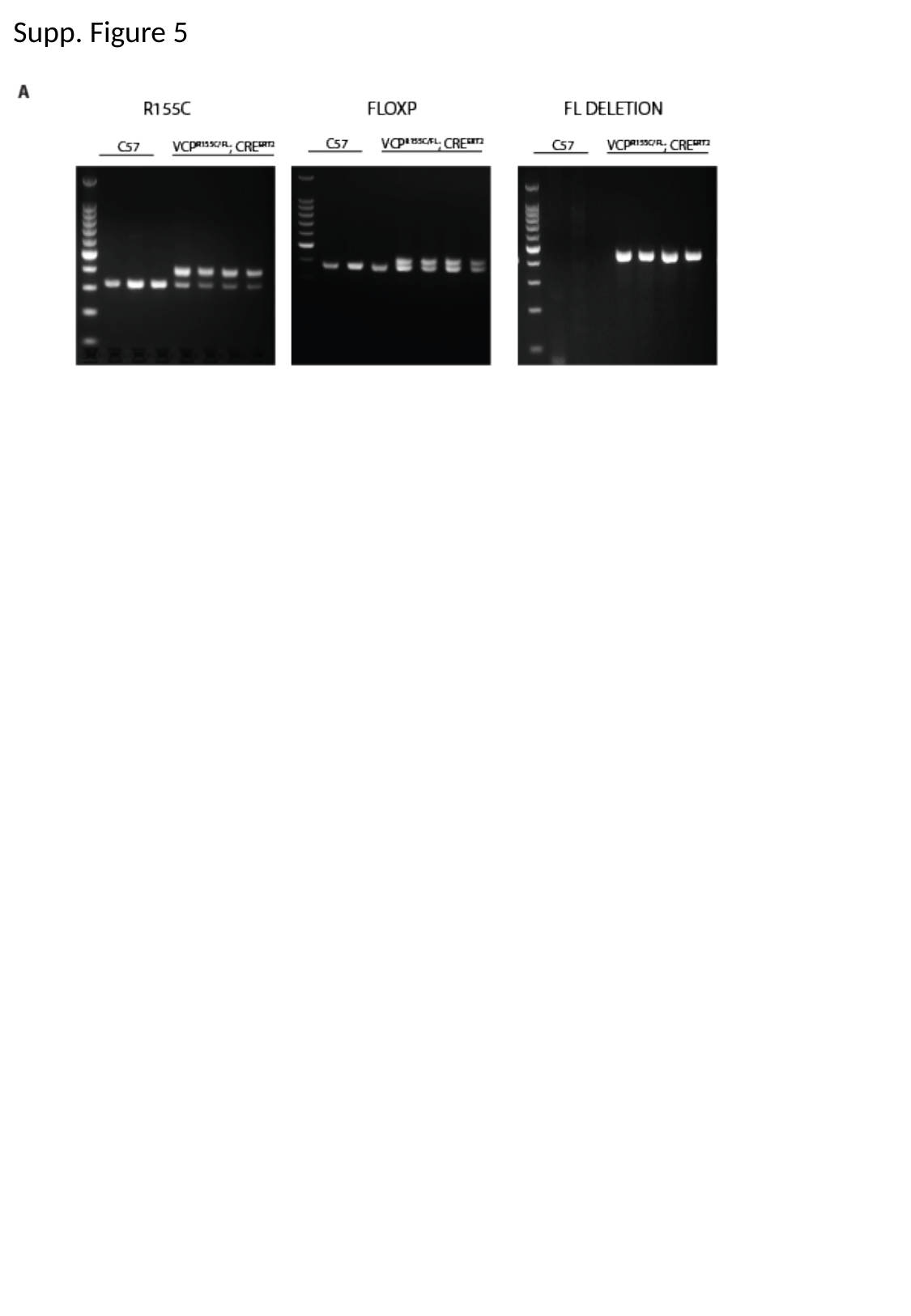

Supp. Figure 5

### Slide 6
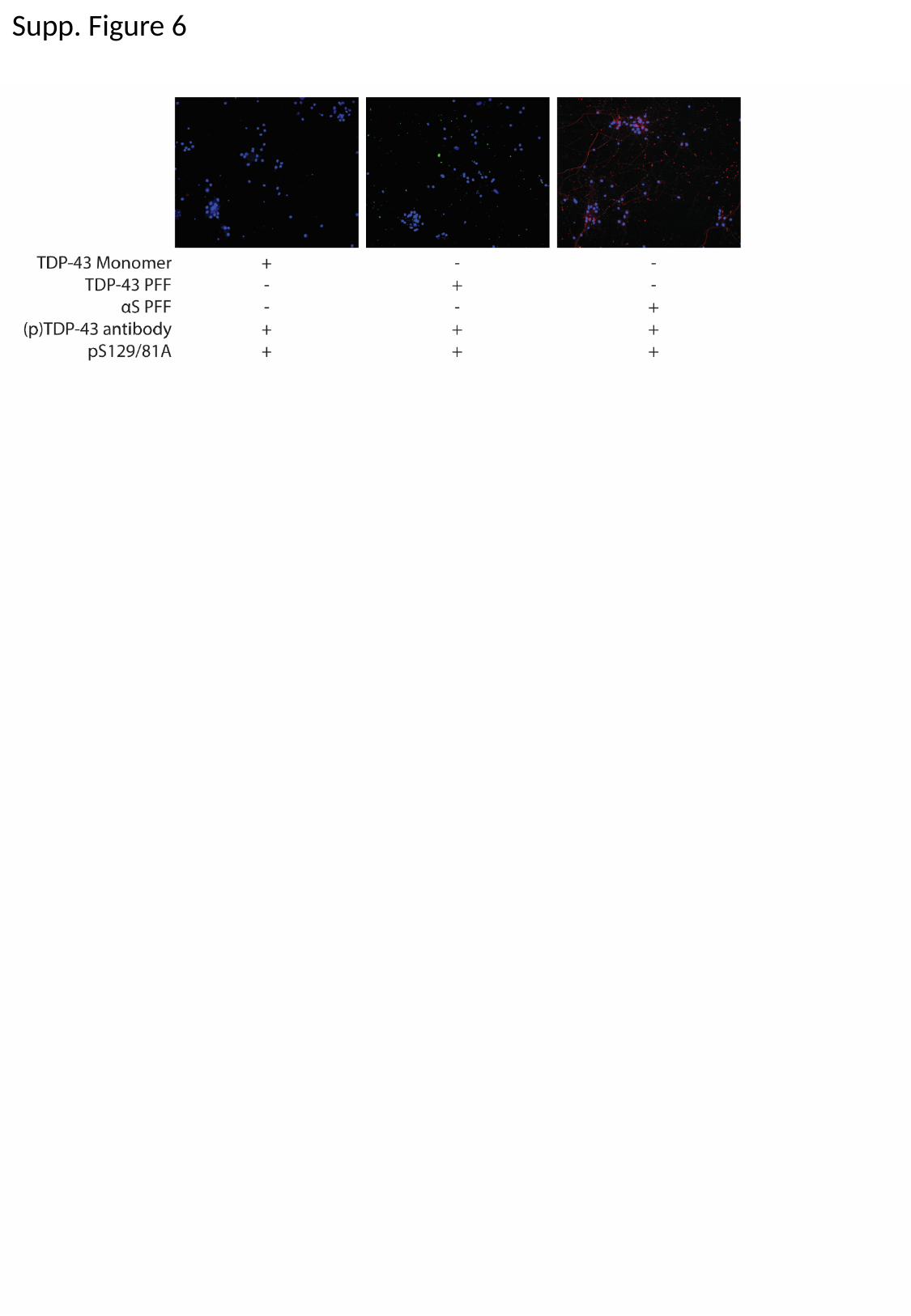

Supp. Figure 6
